# Closed-reference metatranscriptomics enables *in planta* profiling of putative virulence activities in the grapevine trunk-disease complex

**DOI:** 10.1101/099275

**Authors:** Abraham Morales-Cruz, Gabrielle Allenbeck, Rosa Figueroa-Balderas, Vanessa E. Ashworth, Daniel P. Lawrence, Renaud Travadon, Rhonda J. Smith, Kendra Baumgartner, Philippe E. Rolshausen, Dario Cantu

## Abstract

Grapevines, like other perennial crops, are affected by so-called ‘trunk diseases’, which damage the trunk and other woody tissues. Mature grapevines typically contract more than one trunk disease and often multiple grapevine trunk pathogens (GTPs) are recovered from infected tissues. The co-existence of different GTP species in complex and dynamic microbial communities complicates the study of the molecular mechanisms underlying disease development especially under vineyard conditions. The objective of this study was to develop and optimize a community-level transcriptomics (i.e., metatranscriptomics) approach that can monitor simultaneously the virulence activities of multiple GTPs *in planta*. The availability of annotated genomes for the most relevant co-infecting GTPs in diseased grapevine wood provided the unprecedented opportunity to generate a multi-species reference for mapping and quantifying DNA and RNA sequencing reads. We first evaluated popular sequence read mappers using permutations of multiple simulated datasets. Alignment parameters of the selected mapper were optimized to increase the specificity and sensitivity for its application to metagenomics and metatranscriptomics analyses. Initial testing on grapevine wood experimentally inoculated with individual GTPs confirmed the validity of the method. Using naturally-infected field samples expressing a variety of trunk disease symptoms, we show that our approach provides quantitative assessments of species composition as well as genome-wide transcriptional profiling of potential virulence factors, namely cell wall degradation, secondary metabolism and nutrient uptake for all co-infecting GTPs.

## INTRODUCTION

When interacting with their host, plant pathogens do not exist in isolation, but are part of complex and dynamic microbial communities (Allen & Banfield, 2005). Such communities may comprise multiple pathogenic species and other microorganisms with negative, neutral or beneficial interactions while colonizing the same plant organ (Fitt *et al.*, 2006, Turner *et al.*, 2013). Under field conditions, a combination of microbe-microbe and microbe-plant interactions contributes to the development and severity of a disease (Lamichhane & Venturi, 2015). Direct and potentially synergistic relations may develop between the co-infecting microbes, including inter-species signaling (Hogan, 2006, Hosni *et al.*, 2011) and metabolic exchange and complementarity (Zelezniak *et al.,* 2015, Ponomarova & Patil, 2015). Indirectly, by interfering with host immune responses (Nomura *et al.*, 2005), modifying the physicochemical characteristics of the host environment, or killing host cells, pathogens can facilitate host colonization by other microbes, which collectively may exacerbate disease symptoms (Rowe *et al.*, 1985).

Complex diseases that affect woody structures of perennial plants, such as grapevine trunk diseases, are often the result of infection by multiple pathogens, which simultaneously or sequentially colonize the host tissue (Hiscox *et al.*, 2015). All permanent structures of a grapevine can be infected by different fungi that cause distinct trunk diseases (Bertsch *et al.*, 2013). Among these, Botryosphaeria dieback, Esca, Eutypa dieback and Phomopsis dieback are the most common (Bertsch *et al.*, 2013). Trunk pathogens colonize the permanent woody structures of the vine mainly through wounds (Rolshausen *et al.*, 2010). As they spread through the host via a combination of cell wall degradation, toxin secretion and necrosis, damage to the vascular tissues causes the progressive reduction in water and nutrient translocation, and eventually leads to the death of the infected organs and, possibly, of the entire vine (Pouzoulet *et al.*, 2014). Disease symptoms in organs distal to the infected tissue may also develop due to secondary metabolites (*i.e.*, phytotoxins) secreted by the pathogens, as in the case of Eutypa dieback and Esca (Andolfi *et al.*, 2011).

The majority of studies on host-plant interactions in the trunk-disease complex rely on inoculations with a single pathogen (e.g., Camps *et al.*, 2010; Czemmel *et al.*, 2015). Nonetheless, grapevine wood is typically colonized by trunk pathogens and other wood-colonizing fungi, some of which are saprophytes or endophytes (Bruez *et al.*, 2014, Travadon *et al.*, 2016, Péros *et al.*, 1999, Úrbez-Torres *et al.*, 2006, Luque *et al.*, 2009). Such fungi are thought to interact with each other, possibly influencing disease symptoms and severity (Sparapano *et al.*, 2001, Pierron *et al.*, 2016, Whitelaw-Weckert *et al.*, 2013). For example, co-inoculations of the two Esca pathogens *Phaeomoniella chlamydospora* and *Phaeoacremonium minimum* resulted in more severe grapevine decline than single species inoculations. A similar additive interaction was observed in co-inoculations of *Ilyonectria macrodidyma* (one of the causal agents of Black leg) and *Diplodia seriata* (one of the causal agents of Botryosphaeria dieback) (Whitelaw-Weckert *et al.*, 2013). A negative interaction was reported between *Phaeoa*. *minimum* and *Fomitiporia punctata*, an Esca secondary pathogen (Sparapano *et al.*, 2001). These few examples of controlled inoculations help us understand a few idealized scenarios, but no study has attempted so far to dissect the *in planta* activities of the trunk pathogen community under vineyard conditions.

In recent years, next-generation sequencing (NGS)-based approaches have enabled profiling of microbial communities on an unprecedented scale (Vernikos *et al.*, 2015). Methods based on massively parallel sequencing of short DNA fragments amplified by PCR have become broadly applied [i.e., DNA barcoding; (Schoch *et al.*, 2012)]. Genome shotgun sequencing of complex biological samples (i.e., metagenomics) has recently emerged as a more effective approach that overcomes some of the limitations of DNA barcoding (Tyson *et al.*, 2004). Metatranscriptomics, the shotgun sequencing of the community mRNAs, presents an even greater improvement for microbial ecology studies. Unlike methods that target microbial DNA, which cannot differentiate between viable and dead microorganisms, metatranscriptomics targets the metabolically active fraction of the microbiome (Kuske *et al.*, 2015, Fraissinet-Tachet *et al.*, 2014, Bridge & Spooner, 2001).

To date, none of these NGS-based approaches have been used to study the activities of pathogen communities associated with trunk diseases. A key impediment to the application of these methods *in planta* is the low abundance of microbial DNA and RNA relative to host nucleic acids; this results in low sequencing coverage for the entire microbiome, often less than 1% of the total sequencing output (Jones *et al.*, 2014, Blanco-Ulate *et al.*, 2013d, Blanco-Ulate *et al.*, 2015). Insufficient sequencing depth limits the application of metagenomics and metatranscriptomic methods that rely on *de novo* assembly of the metagenome and metatranscriptome (Scholz *et al.*, 2012). The implementation of methods that rely on sequence alignment to references comprising all species potentially associated with a sample is often problematic because of the scarce genomic information available for the species under analysis (Filippidou *et al.*, 2015). To overcome this limitation, we sequenced and assembled the genomes of the most common grapevine trunk pathogens (GTPs): *Eutypa lata, Neofusicoccum parvum, Dip. seriata, Phaeoa. minimum, Phaeom. chlamydospora, and Diaporthe ampelina* (Blanco-Ulate *et al.*, 2013a, Blanco-Ulate *et al.*, 2013b, Blanco-Ulate *et al.*, 2013c, Morales-Cruz *et al.*, 2015). The predicted proteomes of each genome were annotated to obtain a comprehensive catalogue of all potential virulence functions associated with cell wall degradation, secondary metabolism, and nutrient uptake (Morales-Cruz *et al.*, 2015).

Here, we utilized all available genomic references for the most common GTPs to develop a metagenomics and metatranscriptomics approach that relies entirely on NGS read mapping onto a multi-species closed reference. In this work we: (i) developed and optimized a bioinformatic pipeline to precisely align short reads from a biologically complex sample to a reference that comprises multiple species, (ii) tested the bioinformatic pipeline in controlled experiments with artificial inoculation of grape woody stems using single GTP species, (iii) applied the multi-species reference approach to vineyard samples from naturally infected vines showing a variety of trunk disease symptoms, (iv) compared the multi-species reference approach with *de novo* assembly of metatranscriptomics data, and (v) analyzed the expression of putative virulence functions at the GTP community level. Our results demonstrate that mapping-based metatranscriptomics can profile the *in planta* expression of thousands of putative virulence factors of multiple pathogenic species, thereby enabling the study of complex diseases under field conditions at the molecular level.

## RESULTS AND DISCUSSION

### Optimization of a multi-species closed-reference read mapping protocol using simulated datasets

To develop a reference-based metagenomic and metatranscriptomic approach for the *in planta* detection and quantification of trunk pathogens, we first assessed the specificity and sensitivity of popular mapping software using simulated datasets. Unlike traditional plant-microbe assays involving a single pathogen, for which RNA-seq methods are established and widely used (*e.g*, Jones *et al.*, 2014, Blanco-Ulate *et al.*, 2013d, Blanco-Ulate *et al.*, 2015), we had to test software performance with short reads from complex biological samples aligned to a reference that comprises multiple species. The genome sequences of grape and the ten most commonly associated species with grapevine trunk diseases (**Appendix S1 Table S1**) were concatenated into a single multi-species genome reference (total size: 952 Mb; Jaillon *et al.*, 2007, Floudas *et al.*, 2012, Nordberg *et al.*, 2014, Blanco-Ulate *et al.*, 2013a, Blanco-Ulate *et al.*, 2013b, Blanco-Ulate *et al.*, 2013c, Morales-Cruz *et al.*, 2015) to map the metagenomic reads (Fig. 1A). Similarly, transcriptomes of the same species were concatenated to create a multi-species reference of 154,998 protein-coding sequences to map the metatranscriptomic reads. To assess mapping specificity and sensitivity, we generated synthetic Illumina reads from the genomes and transcriptomes of the same organisms that were used as references for read mapping using ART, a next-generation sequencing read simulator (Huang *et al.*, 2012). We generated a total of 40 independent metagenomics and metatranscriptomics simulated datasets by combining variable proportions of the simulated reads from *V. vinifera* (80-100% of the total reads per sample) and from the ten fungal species (0-20% of the total reads per sample; **Appendix S1 Table S2**).

**Fig. 1.**
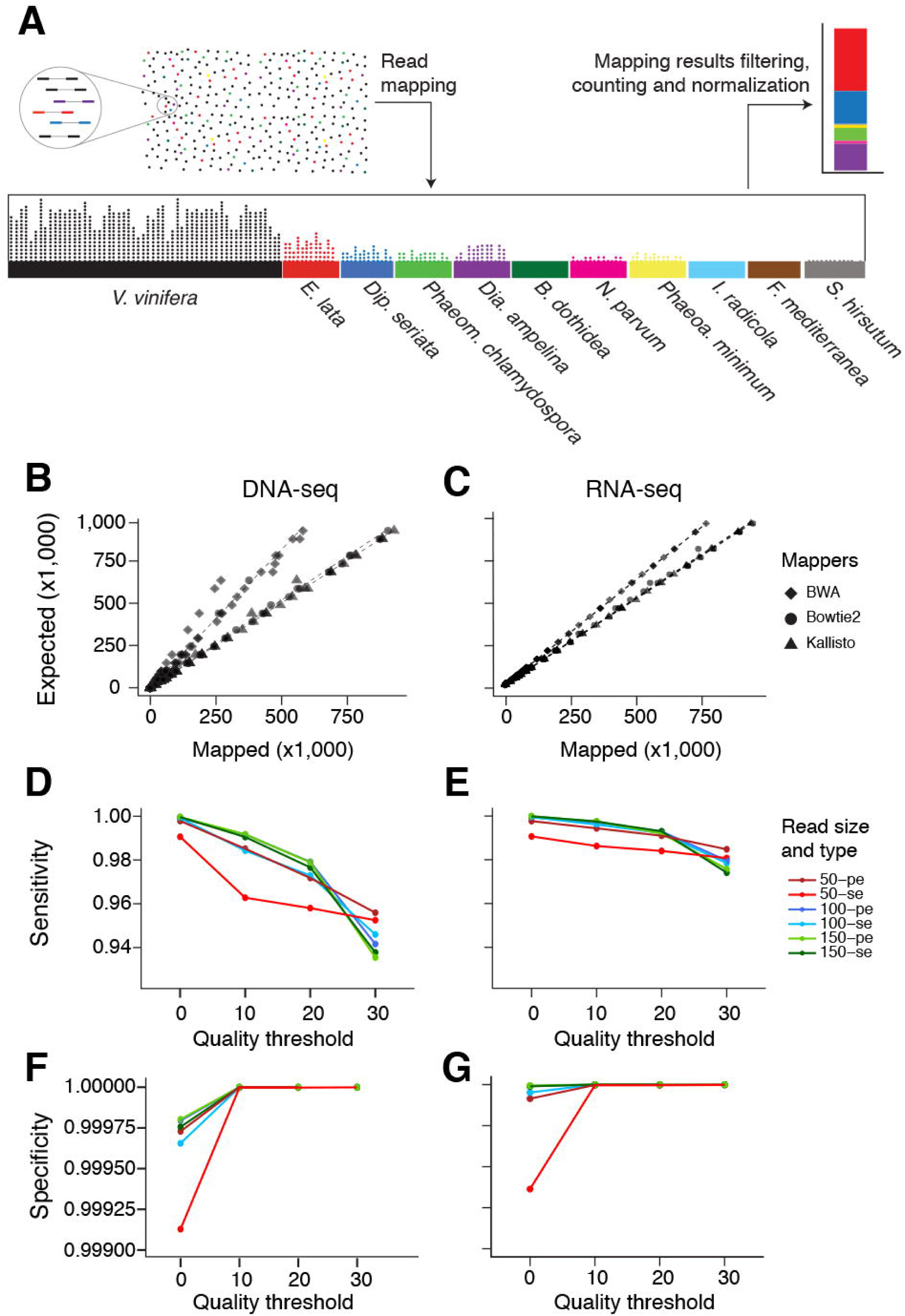
Bioinformatic approach and optimization. (A) Schematic representation of the approach used to quantify sequencing reads for each co-infecting grapevine trunk pathogen species. Dots represent paired-end reads in a given sample, while colors indicate the species from which the reads are derived. Genomic references were combined into a single reference for read mapping. After mapping, the reads are filtered based on mapping quality, counted and normalized. Correlations of the number of simulated short reads with three different short-read mappers and the expected number of reads based on (B) DNAseq and (C) RNAseq data. Evaluation of sensitivity (D, E) and specificity (F,G) when using Bowtie2 with different sequencing parameters and mapping quality thresholds.

The reads of each simulated sample were mapped using three popular short-read aligners: BWA (Li & Durbin, 2009), Bowtie2 (Langmead & Salzberg, 2012) and Kallisto (Bray *et al.*, 2016). Because the taxonomic composition of each of the 20 simulated samples was known, we could determine the sensitivity and specificity of each aligner by comparing expected and observed counts of mapped reads (Fig. 1B&C). Sensitivity (i.e., mapping rate) was high with both Bowtie2 and Kallisto, which aligned 93.5 ± 5.12% and 98.7 ± 1.58% of the expected reads, respectively (Fig. 1A). Mapping rate of BWA was significantly lower (69.89 ± 11.70%; *P* = 0.0019). Rates of non-specific mapping were remarkably low in all tests: Bowtie2 and BWA mapped to the wrong species zero and six (3.97×10^−7^%) reads, respectively, while Kallisto failed to map onto the correct species 11,377 reads (3.12×10^−4^%). Similar patterns of sensitivity and specificity were observed when the simulated metatranscriptomic samples were mapped onto the concatenated multi-species transcriptome (Fig. 1C). We selected Bowtie2 for further optimization because of its high sensitivity and better specificity compared to the other aligners.

Using the simulated datasets, we evaluated the impact on mapping sensitivity and specificity of: (i) increasing threshold of minimum mapping quality (Q0, Q10, Q20, and Q30), (ii) different read lengths (50, 100, and 150 bp) and (iii) different sequencing modes (single *vs*. paired-end; Fig. 1D-G). The weakest sensitivity and specificity were achieved with 50-bp single-end reads and no mapping quality threshold. The maximum specificity was reached at Q10, with little improvement when more stringent parameters were applied. Nonetheless, sensitivity suffered when increasing quality thresholds were applied. We observed a decline of 2.00 ± 1.37% in sensitivity from Q20 to Q30 with only marginal improvement in specificity. Concerning the impact of read length and sequencing mode, the highest specificity was achieved with 150-bp paired-end reads, but with little difference with other combinations. Based on these results, all the analyses described below were carried out using Bowtie2 and a mapping quality cutoff of Q20.

### Controlled inoculations of single pathogen isolates confirm the specificity of the multi-species closed-reference mapping approach

To test the selected and optimized mapping method on infected grapevine samples, we collected and sequenced RNA from vines artificially inoculated with individual GTPs. The woody stems of 12 vines were inoculated with mycelial plugs of either *N. parvum* (isolate UCD646So) or *Phaeoa*. *minimum* (isolate UCR-PA7). From each plant, a stem sample at the inoculation site (±2 cm from the point of inoculation) was processed for RNA extraction and sequencing. In all samples, the inoculated species were detected by metatranscriptomics analysis as the most abundant, confirming the specificity of mapping to a multi-species reference (Fig. 2A&B). On average 90.74 ± 3.90% and 66.61 ± 15.83% of the total fungal reads (i.e., non-*V. vinifera* reads) mapped on *N. parvum* and *Phaeoa*. *minimum*, respectively. Although it was not part of the inoculations, the Esca pathogen *Phaeom*. *chlamydospora* was detected in all assayed plants, albeit at a much lower level than the inoculated pathogens (Fig. 2A&B). On average, *Phaeom*. *chlamydospora* had 15.9 and 4.46 times fewer reads than the most abundant species in the samples inoculated with *Phaeoa*. *minimum* and *N. parvum,* respectively. Given that *Phaeom*. *chlamydospora* has been isolated from asymptomatic grapevine propagation materials (Mugnai *et al.*, 1999, Ferreira *et al.*, 1999, Eskalen *et al.*, 2007), we assume it was present before inoculation. This hypothesis is consistent with its detection in all 12 samples and with little variation in read counts (5,562.08 ± 711.74 reads).

**Fig. 2.**
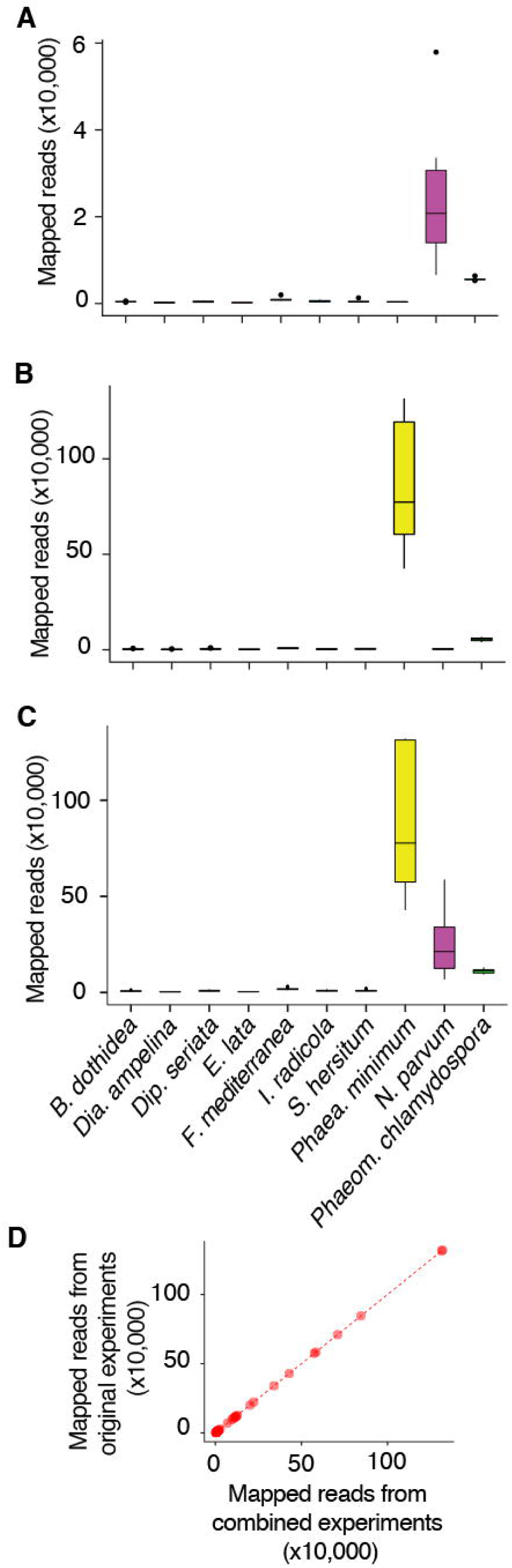
Evaluation of reference-based metatranscriptomics using woody samples inoculated with selected wood pathogens. Grapevine woody stems were inoculated either with *N. parvum* (A) or *Phaeo*. *minimum* (B) and total RNA sequenced. Reads from both experiments were concatenated and mapped together to assess specificity when multiple species are present in the dataset (C, D). Boxplots (A,B,C) show the distribution of read counts mapped onto the different pathogens part on the multi-species reference. Correlation between counts of mapped reads between individual and combined read sets (D).

To further assess specificity when reads from multiple species are mapped, we concatenated the reads obtained from both *N. parvum* and *Phaeoa*. *minimum* inoculations, and mapped the combined reads to the multi-species reference. The simultaneous mapping of the combined dataset led to identical results as when reads were mapped separately (*R* = 1.00; Fig. 2C&D). Both results, the quantitative determination of the inoculated species as the most abundant species and the absolute specificity when mapping reads from multiple species, confirmed that the optimized method is able to assign the reads to the correct species in real experimental data.

### Application of the multi-species closed reference mapping approach to field samples

With the objective of studying GTPs under vineyard conditions, we then applied the multi-species mapping approach to samples taken from mature vines (> 8 years-old) showing a variety of the most common symptoms associated with trunk diseases (**Appendix S1 Table S3**). We collected 28 wood samples from distinct plants with the following combinations of symptoms: Eutypa dieback foliar and wood symptoms (8 samples; Fig. 3A&C), Esca foliar and wood symptoms (8 samples; Fig. 3B&C), wood symptoms and no foliar symptoms (6 samples; Fig. 3C), and apparently healthy plants with no foliar or wood symptoms (6 samples; Fig. 3D). For samples with wood symptoms, the tissue at the margin of the necrotic region was collected (Fig. 3C). DNA and RNA were extracted from each sample, and used for metagenomics (DNA-seq) and metatranscriptomics (RNA-seq) analysis. The same samples were also used for culture-based identification. Fungal colony purification led to the identification of up to two GTPs per sample (**Appendix S2 Fig. S1**), which corresponded to the foliar symptoms. *E. lata* was isolated from all vines with Eutypa dieback symptoms. *Phaeom*. *chlamydospora* and *Phaeoa. minimum*, were recovered from 50% and 38%, respectively, of vines with Esca symptoms. From vines with no foliar symptoms, wood symptoms were associated with the presence of *Phaeom*. *chlamydospora, N. parvum, Dip. seriata* and *Dia. ampelina*, but not *E. lata. Phaeom. chlamydospora, N. parvum, Dip. seriata* were also recovered from vines with Eutypa dieback and Esca symptoms.

**Fig. 3.**
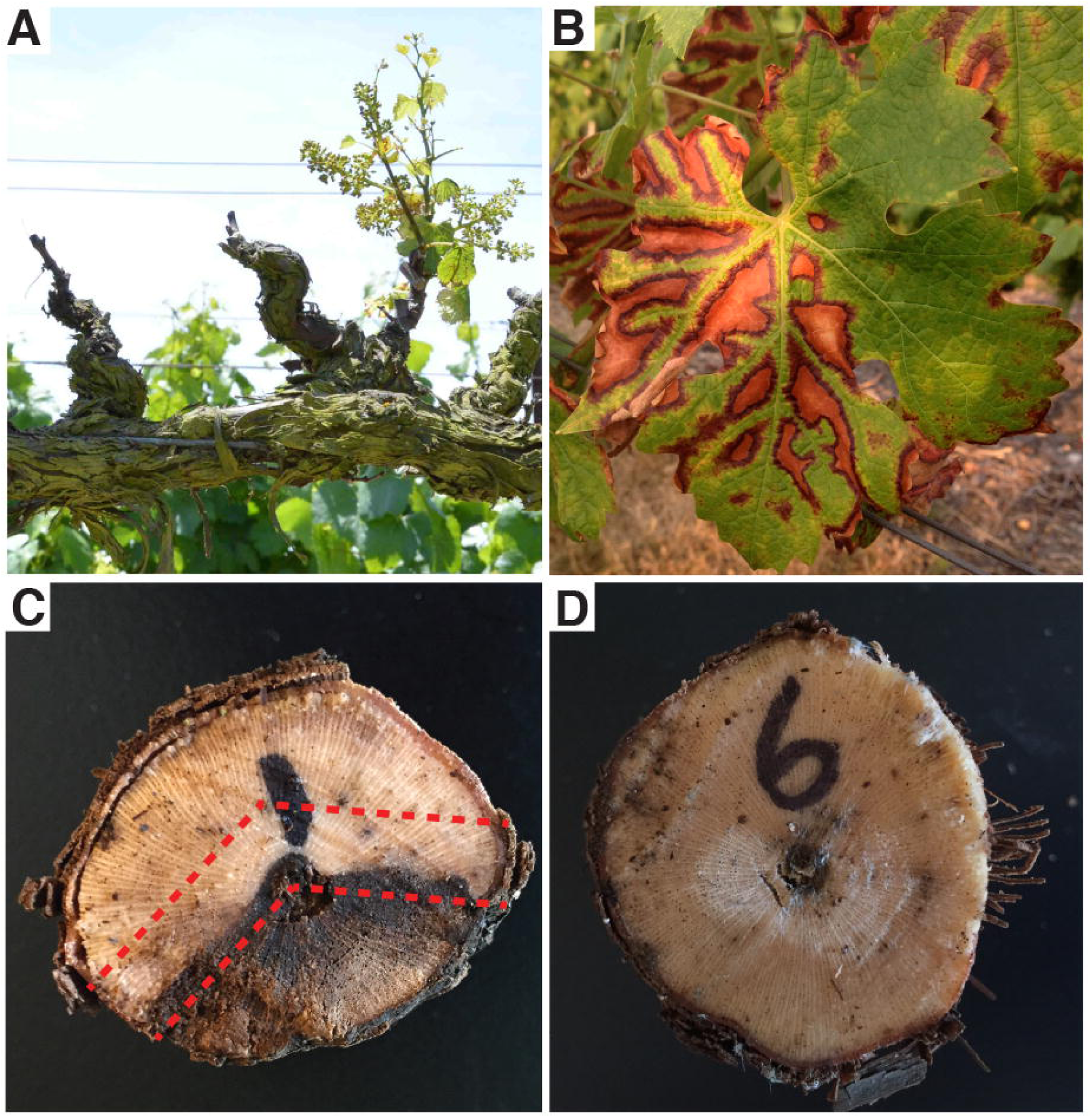
Symptoms of trunk disease infections that were used to classify the field samples used in the study. Foliar symptoms of Eutypa dieback (A) and Esca (B), wood cankers from vines without any foliar symptoms (C), and apparently healthy (i.e. non-symptomatic) wood (D). In panel C, the red dashed line depicts the area of wood at the edge of the canker collected for RNA and DNA extractions.

For metagenomics and metatranscriptomics analysis of the 28 samples, we generated an average of 57.5 ± 16.5 million DNA-seq and RNA-seq reads per sample, of which 52.0 ± 14.0 million reads were retained after quality and size trimming (**Appendix S1 Tables S4 and S5**). A high proportion of the reads were mapped to the multi-species reference (83.17 ± 11.35% of DNA-seq reads and 87.80 ± 4.70% of RNA-seq reads). The majority of the reads were mapped onto *V. vinifera* (95.72 ± 4.45% and 99.99 ± 0.01%, from plants with and without foliar symptoms, respectively). A greater fraction of reads mapped onto the GTP references when samples were taken from plants with foliar symptoms (4.28 ± 4.45%) compared to those without (0.01 ± 0.01%; Fig. 4A). The reads that did not map to the multi-species reference may derive from organisms that are not included in the reference, as suggested by the taxonomic analysis of *de novo* assembled contigs described below. To quantify the taxonomic composition of the samples, we removed the reads that mapped onto the grape genome and normalized the counts to account for both uneven sequencing throughput and differences in genome or transcriptome sizes between species. Species composition profiles were then constructed using the normalized counts (Fig. 4B&C). The detected species and their relative abundances matched what we expected based on the visible foliar symptoms and what we know about their aetiology. *E. lata* was the predominant species in the samples from vines with Eutypa dieback symptoms with 83.05 ± 10.68% of the reads assigned to the GTP. In all of these samples, however, *Phaeom. chlamydospora* (7.49 ± 8.47%) and *Dip. Seriata* (6.69 ± 3.93%) were also detected at high levels. Overall, results of metagenomics and metatranscriptomics revealed a greater species complexity than suggested by fungal isolation. This may be due to the fact that culture-based identification can favor fungi that grow rapidly (Bridge & Spooner, 2001). The advantage of applying a sequencing method *in planta* was particularly evident in samples from vines with Esca foliar and wood symptoms and vines with no foliar symptoms. In the former, both Esca pathogens were consistently detected as the predominant species in all samples together with smaller amounts of *Dia*. *ampelina*. Vines with wood symptoms and no foliar symptoms had by far the most complex and variable composition, ranging from mostly *Dia*. *ampelina* to mostly *Phaeom*. *chlamydospora*, or different proportions of these two species in combination with *Dip*. *seriata*. Much lower fungal counts, mostly associated with *Phaeom*. *chlamydospora* and *Phaeoa*. *minimum*, were detected in apparently healthy (i.e. no foliar nor wood symptoms of trunk diseases) plants. Although both of these species are reported from apparently healthy plants (Pancher *et al.*, 2012, Bruez *et al.*, 2014, Casieri *et al.*, 2009, Hofstetter *et al.*, 2012), the quantitative nature of the mapping method determined the symptomatic samples had close to 197 times more reads on average than the apparently healthy plants (Fig. 4A). These results clearly show that even if potentially pathogenic species can be detected in healthy samples, their abundance is significantly greater in plants with disease symptoms. Furthermore, this suggests that pathogen detection methods aiming to differentiate between the early and late stages of infection should be quantitative.

**Fig. 4.**
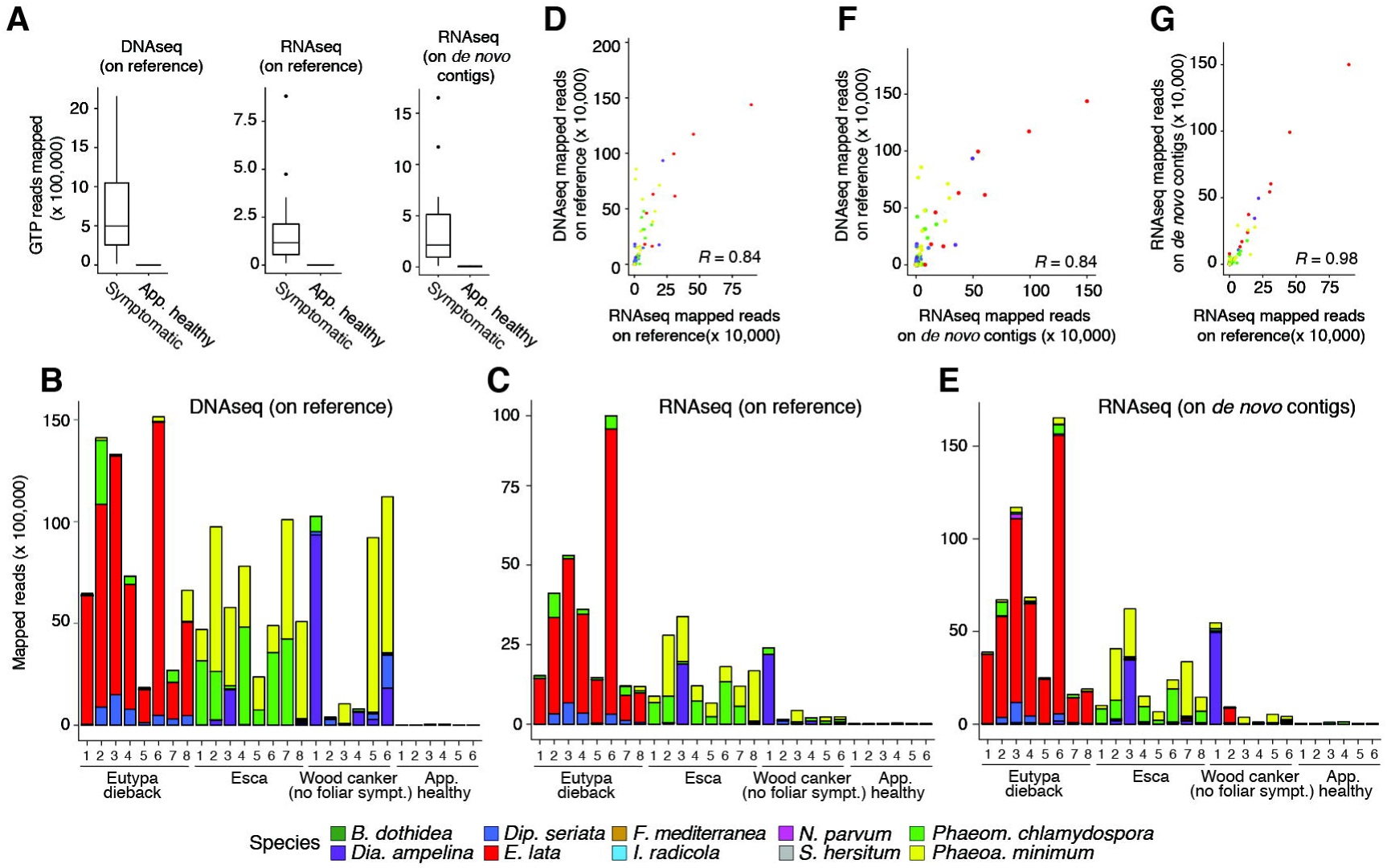
Metagenomics and metatranscriptomics analyses of field samples. (A) Boxplots show distribution of counts of mapped reads onto the multi-species genome and transcriptome or *de novo* assembled contigs. Stacked bars show counts of mapped reads per each trunk pathogen species in the reference from DNA-seq (B) and RNA-seq (C) analyses. (D) Scatterplot shows the correlation of counts per species determined using DNA-seq and RNA-seq. (E) Stacked bars show counts of mapped reads on *de novo* assembled contigs assigned to the different trunk pathogen species. (F,G) Scatterplots show the correlation of counts per species determined using reference-based approaches (DNA-seq in F, RNA-seq in G) and mapping on *de novo* assembled contigs.

Species profiles determined by metagenomics showed a significant linear correlation with the results of metatranscriptomics (R = 0.84; *P* = 2.2 e^−16^; Fig. 4F). This correlation is an independent validation of the relative amount of each species using the two different methods. The two methods in fact do not only involve two different types of nucleic acids, but also different extraction methods, library preparation, and mapping references. Some of the difference between results of DNA-seq and RNA-seq mapping values per species may be due the different metabolic activities of the fungi present in the sample at the time of collection. This was particularly evident in the samples with wood symptoms and no foliar symptoms, in which fungal DNA and RNA quantities had a lower correlation.

### Metatranscriptomics based on *de novo* assembly validates the multi-species closed reference mapping approach

A potential limitation of the reference-based approach described above is that mapping is restricted to a closed reference. This may force reads to align to the concatenated transcriptomes even if they derive from species not included in the multi-species reference. To evaluate the impact of restricting the mapping of the metatranscriptomics reads to a closed reference, we compared results of the multi-species closed reference approach with results of read mapping onto *de novo* assembled contigs (**Appendix S2** Fig. 2). Transcripts of the 28 samples were assembled *de novo* using the quality-filtered RNA-seq reads. As a pre-filtering step, we excluded from the assembly process the reads that mapped to the transcriptome of *V. vinifera*. A total of 662 million reads that did not map to the *V. vinifera* reference were assembled into 2.3 million contigs using MEGAHIT (Li *et al.*, 2015). An average of 80.9 ± 18.3 million contigs per sample were obtained. We then assigned contigs to taxonomic groups by homology to the NCBI protein RefSeq database using BLASTx (e-value < 1e^−6^). To avoid false taxonomic assignments due to missing information in the database, we limited the taxonomic assignments to the genus level. Despite the pre-filtering steps, an average of 85.01 ± 11.87% of the contigs were still assigned to *V. vinifera* proteins in RefSeq. A total of 167 fungal genera were detected across the 28 samples (**Appendix S1 Table S6**), including nine of the ten genera of the multi-species reference (all except *Botryosphaeria*). Asymptomatic samples showed significantly smaller amounts of fungal contigs (0.16 ± 0.08% of the total contigs) compared to the symptomatic samples (10.99 ± 6.50% of the total contigs; P=1.82e^−6^). On average 46.64 ± 25.99% of the fungal contigs were assigned to the GTP present in the closed-reference. Overall, the assembled GTP transcripts were shorter (663.35 ± 27.50 bp) than the predicted CDS in the GTP genomes (1,422.30 ± 1,076.37 bp), indicating fragmentation and potential redundancy of the fungal transcripts in *de novo* assembled contigs (Fig. 5A). The rest of the contigs (53.36 ± 25.99% on average per sample) were assigned to 158 non-GTP taxa (**Appendices S1 Table S6 and S2** Fig. 3).

**Fig. 5.**
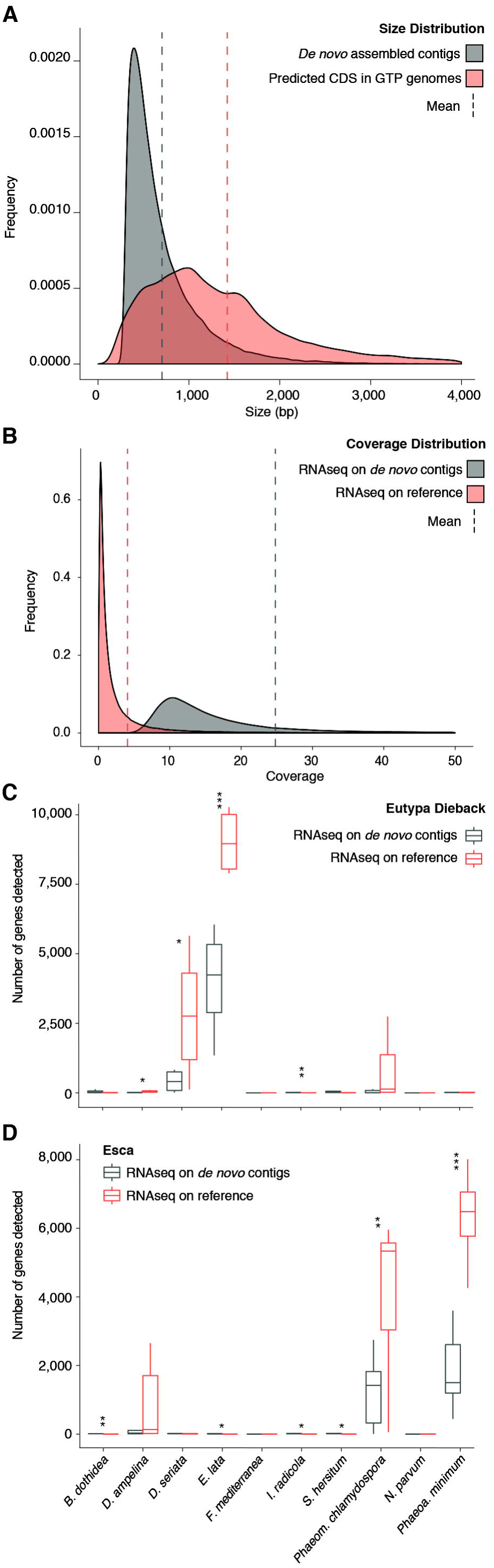
Evaluation of the metatranscriptomics using the reference-based method or *de novo* assembly. Size (A) and RNA-seq read coverage (B) distribution of contigs *de novo* assembled and the protein-coding sequences (CDS) predicted from the genomic references. (C,D) Boxplots summarizing the distribution of number of detected protein-coding genes assigned to each pathogen species in samples with Eutypa dieback (C) and Esca (D) symptoms. Asterisks indicate significant differences in means (\**P* ≤ 0.05, \*\**P* ≤ 0.01, *** *P* ≤ 0.001).

The abundance (i.e. expression level) of each *de novo* assembled contig was measured by mapping the metatranscriptomics reads onto the contigs. An average of 0.57 ± 0.78 million reads per sample were mapped to *de novo* contigs assigned to GTPs. The median mapping coverage per contig was 13.8x (Fig. 5B), approximately three-fold higher than the median coverage obtained by the closed reference-based approach. The overall higher coverage of mapping on *de novo*-assembled contigs suggests that transcripts with higher expression levels were preferentially assembled by MEGAHIT. This hypothesis is also supported by the greater number of genes per sample detected by the closed reference-based approach (8,711.46 ± 6,055.12 CDS/sample) compared to the number of mapped *de novo* contigs (7,522.36 ± 6,889.68 contigs/sample). The smaller number of contigs compared to the total CDS used in the reference-based approach may have also contributed to increase the median coverage. To estimate the difference in the number of genes detected by the two metatranscriptomics approaches, we assigned homology between the *de novo* assembled contigs and the predicted CDS that compose the multi-species reference using BLASTn (e-value < 1e-6; Fig. 5). In most cases, the reference based-approach detected a significantly larger number of CDS than the *de novo* assembly approach (Fig. 5). Based on these results we can conclude that the *de novo* assembly approach yielded a narrower representation of the metatranscriptome limited to those transcripts with enough coverage to allow assembly.

The taxonomic profiles obtained using the *de novo* assembly were very similar to those obtained using the reference-based metatranscriptomic (*R*=0.98; *P*=2.2e^−16^; Fig. 4E&F) and metagenomic (*R*=0.84; *P*=2.2e^−16^; Fig. 4G) data. These results confirmed that the multi-species-reference approach did not lead to false taxonomic assignments at least at the genus level due to the closed-reference. On average, 53.33 ± 19.58% of the reads mapped to contigs from GTPs. *Phaeomoniella, Eutypa, Diaporthe* and *Phaeoacremonium* were often the predominant genera (**Appendix S2** Fig. 3). For example, the genus *Phaeomoniella* accounted for up to 93.1% of reads in sample AH2, *Eutypa* for 64.9% in sample ED7, *Diaporthe* for 60.6% in sample WC1 and *Phaeoacremonium* for 51.6% in sample WC3. Most of the remaining 158 non-GTP genera contributed individually to less than 5% of the total number of reads per sample and collectively to 35.21 ± 12.55%. The few genera that contributed to more than 5% in at least one sample (**Appendix S2** Fig. 3) included known plant pathogens or endophytes, such as *Aspergillus, Pestalotiopsis, Leptosphaeria, Stagonospora, Bipolaris, Pyrenophora*, and *Setosphaeria* (Nierman *et al.*, 2005, Kumar *et al.*, 2002, Li & Strobel, 2001, Howlett *et al.*, 2001, Eyal, 1999, Lamari & Bernier, 1989, Perkins & Pedersen, 1987, Munkvold & Marois, 1995) as well as other ubiquitous fungi (Longcore *et al.*, 1999, Feng *et al.*, 2014, Corte *et al.*, 2015). Overall, the greater contribution of GTP genera to the total read count per sample suggests a dominant role of these genera in the fungal community.

In conclusion, while the *de novo* assembly approach validated the results of the reference-based approach, it provided a narrower and more fragmented representation of the GTP community’s metatranscriptome. These limitations, combined with a more complex pipeline (**Appendix S2** Fig. 2), longer processing time, and more intensive computational requirements, provides further support to the reference-based approach as the more effective method for profiling transcriptional activities *in planta* of fungal community of known composition.

### Multi-species closed reference-based metatranscriptomics allows the *in planta* profiling of virulence function expression

Phytotoxic metabolites and cell wall degrading enzymes are considered key pathogenicity and virulence factors underlying trunk disease development (Andolfi *et al.*, 2011, Bertsch *et al.*, 2013, Rolshausen *et al.*, 2008). We previously annotated all predicted protein-coding genes of all GTPs’ genomes included in the present work (Morales-Cruz *et al.*, 2015). Functional annotations were assigned based on the presence of conserved domains as well as homology to proteins from relevant specialized databases focused on potential virulence factors involved in primary and secondary cell wall decomposition (i.e. carbohydrate active enzymes, CAZymes), secondary metabolism (i.e. cytochrome P450s, biosynthetic gene clusters) and nutrient uptake (i.e. transporters). The first study that profiled the transcriptional activities of these virulence factors during experimental infections with *N. parvum* of grapevine woody stems revealed that physically clustered genes coding for putative virulence functions share common regulatory sequences and are induced depending on substrate or stage of plant infection (Massonnet *et al.*, 2016). In this work, we utilized the optimized reference-based metatranscriptomics approach to obtain information of *in planta* expression of virulence genes when multiple GTPs are co-infecting the same host under vineyard conditions.

We first extracted the normalized mapping counts of all genes assigned to any of the five broad categories of putative virulence factors (CAZymes, cytochrome P450s, peroxidases, genes belonging to biosynthetic gene clusters, and transporters; **Appendix S1 Table S7**). A total of 530 functions were identified from 12,951 different genes across all samples and GTP species. CAZymes and transporters were the most expressed functions followed by secondary metabolism, cytochrome P450s and peroxidases in all pathogens (Fig. 6A). Expression differences between different potential virulence factors became clear when we analyzed the expression patterns of specific functions. A Principal Component Analysis (PCA) based on expression data of each virulence function clearly separated samples based on the type of disease symptoms they were associated with (Fig. 6B). Separations due to the two principal components were confirmed by a Partial Least Squares Discriminant analysis (PLS-DA, R^2^Y= 0.988, Q^2^Y=0.903, **Appendix S2** Fig. 4). These results suggest that pathogen species associated with the same disease, even if phylogenetically distant, as in the case of Esca pathogens, activate similar virulence functions at comparable expression levels.

**Fig. 6.**
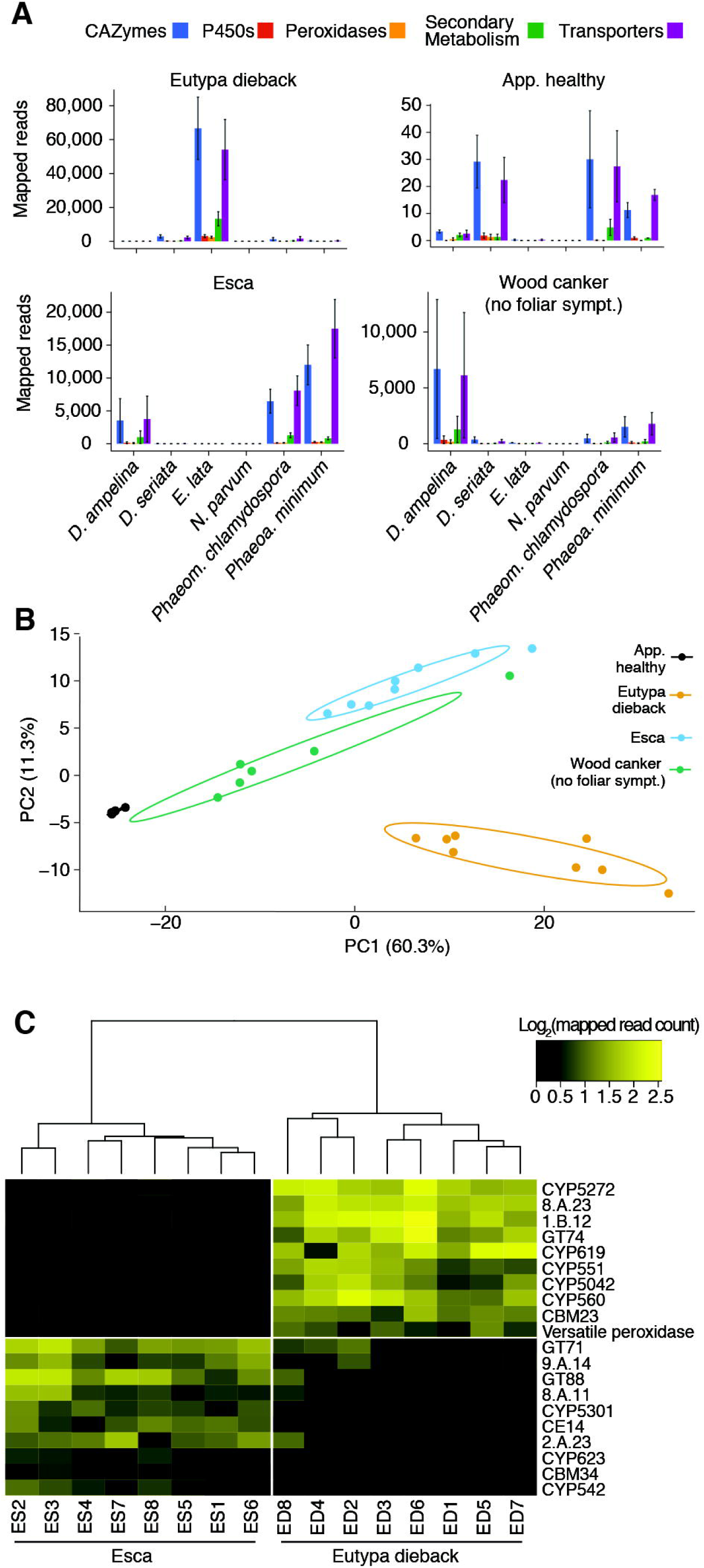
Virulence function expression based on metatranscriptomics analysis. (A) Barplots showing the expression levels measured as normalized read counts of genes grouped by broad functional categories with potential virulence activity for each trunk pathogen species. Principal Component Analysis using as input the sum of reads for each samples that mapped on all genes the share the same Pfam annotation (B). (C) Heatmap showing the expression levels of specific functions identified as major PLS-DA loading separating samples with Esca and Eutypa dieback symptoms.

To explore some of the functions most distinctively expressed between samples with Esca and Eutypa dieback symptoms we extracted the major loadings separating the two groups based on the PLS-DA. For Eutypa dieback samples the major loadings comprised genes potentially associated with cell wall degradation, secondary metabolism and toxin secretion (Fig. 6C): a Versatile Peroxidase (VP), two CAZymes (GT74 and CMB23), five cytochrome P450s, and two transporters. VPs were shown to degrade lignin in other fungi (Camarero *et al.*, 1999, Ruiz-Dueñas *et al.*, 2009). GT74 and CMB23 CAZymes have α-1,2-L-fucosyltransferase and mannan-binding function and, therefore, may both be potentially involved in the host cell wall manipulation (Stoll *et al.*, 2000, Perrin *et al.*, 1999). The identified cytochrome P450s included CYP560, CYP551, and CYP619 whose activities have been associated to fungal secondary metabolism (Moktali, 2013, Artigot *et al.*, 2009). In *Aspergillus clavatus,* CYP619 was shown to participate in the biosynthesis of the toxin patulin (Artigot *et al.*, 2009). One of the two transporters identified belonged to the Autotransporter-1 Family (AT-1, code 1.B.12), which may facilitate the secretion of toxins as seen in bacteria (Tang & Saier, 2014).

Among the functions that more strongly contributed to separate Esca samples, we also found interesting putative virulence functions. These included: a transporter from the family Immunophilin-like Prolyl:Peptidyl Isomerase Regulator (I-PPI, code 9.A.14), which has been extensively studied for its role in exclusion of antifungal drugs in yeast (Arevalo-Rodriguez *et al.*, 2004); CYP542, which has been associated with secondary metabolism (Moktali, 2013) and CYP655, similar to the polyketide synthase (PKS)-NRPS hybrid responsible for the biosynthesis of the tenellin toxin in *Beauveria bassiana* (Xiao *et al.*, 2012). CAZymes GT71 and CBM34 with potential *α*-mannosyltransferase and granular starch-binding activities, respectively, were also characteristics of the Esca samples and may be associated with the degradation of host cell wall or storage polysaccharides (Crucello *et al.*, 2015, Machovič & Janeček, 2006).

## CONCLUSIONS

In this study, we show that existing mapping software can be optimized and applied to study fungal communities associated with grapevine trunk diseases. We demonstrate that a closed reference that includes the most important species associated with a complex biological system can effectively overcome the limitation of using *de novo* assembled contigs and take advantage of the existing gene models and functional annotations. This multi-species reference can be expanded as more genomes of fungi inhabiting grapevines are sequenced. The highly specific and quantitative nature of the mapping approach can find useful application for disease diagnostics both in production vineyards and nurseries, particularly. This metatranscriptomic approach not only can be applied to field samples, as shown here, but also to controlled co-inoculations with different combinations of multiple pathogens under the same environmental conditions to determine the patterns of gene expression as disease symptoms develop in the different organs. Community-level transcriptional analysis integrated with chemical analysis of secreted toxins will help determine the relative contribution of each co-infecting agent to the development of different types of trunk diseases.

## EXPERIMENTAL PROCEDURES

### Optimization of read mapping onto a multi-species closed reference

The references of the genome and transcriptome references available of grapevine trunk associated fungi were concatenated and used as a multi-species reference. This reference was used as input in the ART simulator (Huang *et al.*, 2012) to generate *in silico* reads with the characteristics of reads produced by an Illumina HiSeq 2500 sequencer. From the simulated reads, a total of 20 synthetic samples were produced with different proportions of each species to test the mapping method under different scenarios. Reads were mapped with Bowtie2, BWA and Kallisto with different size lengths (50, 100 and 150 bp) and mapping quality filters (Q value 0, 10, 20, 30). Each combination was evaluated by sensitivity and specificity values.

### Controlled inoculation of *Phaeoa*. *minimum* and *N. parvum*

In March 2014, one-year-old dormant cuttings of *Vitis vinifera* ‘Cabernet Sauvignon’ clone 29 and ‘Merlot’ clone 15, were cut to uniform length (~10 cm) containing two nodes. Cuttings were surface-sterilized in 1% sodium hypochlorite for 15 min and soaked in water overnight and stored within hermetic plastic bags in a cold room (2°C). In April 2014, on the day of inoculations, each cutting was wounded at approximately 3 cm below the uppermost node with a 3-mm cork borer. A 3-mm mycelial plug from a 3-day-old culture of *Neofusicoccum parvum* isolate UCD646So and from a 10-day-old culture of *Phaeoacremonium minimum* (*syn. Togninia minima*) isolate UCR-PA7 was inserted into the wound, and sealed with Vaseline (Unilever, Rotterdam, London, UK) and Parafilm (Bemis Co., Neenah, WI, USA). Non-inoculated controls were wounded and ‘mock-inoculated’ with an agar plug from a sterile petri plate of Potato Dextrose Agar (PDA; Difco laboratories, Detroit, MI). After inoculation, cuttings were submerged in melted paraffin wax within 4 cm of the roots to prevent moisture loss as roots and shoots formed. Cuttings were potted in a mix of perlite and vermiculite (1:1) into aerated plant bands (5 ×5 × 20 cm; Monarch Manufacturing Inc., Salida, CO) held into plastic trays with partially opened bottom allowing appropriate aeration and drainage. Inoculated plants were supplied with bottom heat to enhance rooting process by placing plastic containers onto heating pads; the minimal rooting temperature was maintained above 24°C at night. Plants were incubated in the greenhouse [natural sunlight photoperiod, 25 ± 1°C (day), 18 ± 3°C (night)]. Two replicate experiments were conducted over two successive weeks in two distinct greenhouses. In each experiment, plants were arranged in a completely randomized design, with six inoculated and six mock-inoculated plants per cultivar.

Because the pathogens colonize wood at different rates, infection was confirmed after inoculation at 2 and 10 months post inoculation for *N. parvum* and *P. minimum*, respectively, by observation of an internal lesion (i.e., wood discoloration surrounding the inoculation site). First, the green shoots, roots, and bark of each plant were removed and discarded, and the woody stems were surface sterilized in 1% sodium hypochlorite for 2 min and rinsed with deionized water. Then the shoot was cut longitudinally, to reveal the lesion. To confirm the pathogens were responsible for the lesion’s presence in inoculated plants, and its absence from non-inoculated plants, pathogen recovery was attempted by cutting ten pieces (2 × 5 × 5 mm) of wood from the distal margin of the lesions followed by surface disinfestation in 0.6% sodium hypochlorite (pH 7.2) for 30 s, rinsed twice for 30 s in sterile deionized water, plated on PDA amended with tetracycline (5 mg/ml), and incubated in the dark at 22°C for 14 to 21 days. Wood cylinders app. 2 cm in length were collected with flame-sterilized forcepss right beneath the inoculation site, placed in liquid nitrogen and stored at −80°C until processed for nucleic acid extractions.

### Symptomatic and non-symptomatic sample collection from mature vines

In 2014 and 2015, 28 wood samples were collected from vineyards affected by trunk diseases. One wood sample was collected per vine so that a total of 28 vines were sampled. The vineyards were located in different California grape production areas and included wine grapes cv. Pinot Meunier (14 samples) located in Sonoma County, and table grapes cvs. Flame (7 samples), Dovine (1 sample), Thompson Seedless (3 samples) and Crimson (3 samples) located in Fresno County. Vines were selected based on wood and/or foliar disease symptom expression as previously described (Gubler *et al.*, 2015; Rolshausen *et al.*, 2015). Each collected sample was cut into two wood pieces, half was used for the culture-based fungal identification while the other half was submerged in liquid nitrogen and stored at −80°C for nucleic acid extraction.

### Culture-based identification method

Fungal isolates were recovered from woody necrotic areas (i.e., showing signs of streaking, canker, browning) on PDA amended with tetracycline (100 ppm), with two plates per sample. Wood chips (approx. 3 × 3 × 3 mm in size) were removed from the necrotic areas with a sterile blade and disinfested in 10% bleach (sodium hypochlorite) for 2 min. and rinsed twice in distilled water for 2 min. Plates were incubated at room temperature in the dark and inspected several times per week for two weeks. Fungal isolates with culture morphologies typical of *Botryosphaeria, Diaporthe, Diplodia, Eutypa, Neofusioccum, Phaeoacremonium* and *Phaeomoniella*, and were hyphal-tip purified and transferred to PDA. We recovered a total of 6 isolates from wood samples that were further identified by ITS rDNA sequencing. DNA was extracted from mycelium scraped from the surface of 14-day-old cultures grown at room temperature (DNeasy^®^ Plant kit Qiagen, Valencia, CA, USA), following manufacturer instructions. The nuclear loci rDNA Internal Transcribed Spacer (ITS1/5.8S/ITS2) were amplified using PCR primers ITS1 and ITS4 (White *et al.*, 1990). PCR was performed with cycling parameters of one cycle at 94°C for 5 min, 35 cycles at 94°C for 1 min, 58°C for 1 min and 72°C for 1.5 min, and a final elongation step at 72°C for 5 min. PCR products were sequenced in both forward and reverse directions (Genomic Core Sequencing Facility, University of California, Riverside). BLASTn searches of GenBank identified homologous sequences with high identity.

### Nucleic acid extraction and sequencing library preparation

Frozen wood pieces were ground using the TissueLyser II (Qiagen, Valencia, CA, USA) with stainless steel jars frozen in liquid nitrogen into a fine powder. Part of the ground tissue was used to extract DNA using a modified version of the protocol described in Stoffel *et al.* (2012) with 200 mg of initial ground tissue. The DNA was assayed for concentration, purity and integrity with Qubit (Life Technologies, Eugene, OR, USA), Nanodrop 2000 (Thermo Scientific, Waltham, MA, USA) and an agarose 1X gel, respectively. High quality DNA was fragmented by sonication using the Bioruptor. Fragmented DNA was used as template for library preparation using a KAPA Biosystems Illumina kit. Size selection after ligating adapters was done with the eGel system (Invitrogen, Carlsbad, CA, USA). Final libraries were sequenced in the Illumina HiSeq3000 as 150 bp paired end. Another aliquot of the ground wood tissue was used to extract RNA using the protocol described in Blanco-Ulate *et al.* (2013d). RNA concentration and purity was evaluated similarly than the DNA, while integrity was evaluated with an 2% agarose gel. The RNA was used as input for the Illumina TruSeq Kit for library preparation. Final libraries were sequenced in the Illumina HiSeq300 as 150 bp paired end.

### *De novo* assembly of metatrascriptomic data and taxonomy assignment

Quality filtered reads of the metatranscriptomics data were mapped to the *V. vinifera* genome reference using Bowtie2 (as described above). The reads that did not map to the *V. vinifera* genome were used as input for the assembler MEGAHIT (Li *et al.*, 2015). The software was run on the meta-sensitive mode and only contigs larger than 300 bp were kept from the contigs generated. The reads used to generate the contigs were mapped with Bowtie2 onto the assembled contigs to determine the contig expression levels. Contigs with counts < 10 reads and/or that that aligned (e-value < 10^−6^) to *V. vinifera* were removed. The RefSeq protein database of fungi and *V. vinifera* was used as database to assign taxonomic memberships to the *de novo* assembled contigs using BLASTx (e-value < 10^−6^). The taxonomy of the protein with best hit of each contig was extracted and used to assign a genus to the aligned contig. Finally, the number of reads per taxonomic unit was used to create an abundance profile based on the number of reads.

## ACKNOWLEDGMENTS

This work was supported by the American Vineyard Foundation (grants 2014-1798 and 2015-1798) and by the USDA, National Institute of Food and Agriculture, Specialty Crop Research Initiative (grant 2012-51181-19954). AMC was partially supported also by The Horace O. Lanza Scholarship, The Pearl & Albert J. Winkler Scholarship in Viticulture and the André Tchelistcheff and Dr. Richard Peterson Scholarship.

## Accession numbers

BioProject ID: PRJNA352065

SRA accession number: SRP092409

## Supporting information legends

### Appendix 1. Supplemental Tables

**Table S1.** Genomes and transcriptomes used to generate the multi-species reference

**Table S2.** Composition of simulated samples used for testing and optimization of read mapping

**Table S3.** Description of the field samples used in the study including the culture-based diagnosis of the associated trunk diseases

**Table S4.** DNAseq sequencing and mapping metrics of the field samples

**Table S5.** RNAseq sequencing and mapping metrics of the field samples

**Table S6.** Fungal genera detected in each sample based on homology to peptides in the NCBI protein RefSeq database (BLASTx; e-value < 1e-6)

**Table S7.** Read counts of RNAseq per function detected in each sample

### Appendix 2. Supplemental Figures

**Fig. S1:** Taxonomy of the grapevine trunk pathogens isolated from the field samples.

**Fig. S2:** Pipeline used for taxonomic assignment of *de novo*-assembled RNAseq contigs.

**Fig. S3:** Barplot showing the abundance of the genera detected by mapping RNAseq data onto the *de novo* assembled contigs.

**Fig. S4:** Description of the PLS-DA model using the R package “ropls”.

